# AncesTree: an interactive immunoglobulin lineage tree visualizer

**DOI:** 10.1101/2020.02.17.952465

**Authors:** Mathilde Foglierini, Leontios Pappas, Antonio Lanzavecchia, Davide Corti, Laurent Perez

## Abstract

High-throughput sequencing of human immunoglobulin genes allows analysis of antibody repertoires and the reconstruction of clonal lineage evolution. Phylip, an algorithm that has been originally developed for applications in ecology and macroevolution, can also be used for the phylogenic reconstruction of antibodies maturation pathway. The study of antibodies (Abs) affinity maturation is of specific interest to understand the generation of Abs with high affinity or broadly neutralizing activities. Phylogenic analysis enables the identification of the key somatic mutations required to achieve optimal antigen binding. To complement Phylip algorithm, we developed AncesTree, a graphic user interface (GUI) that aims to give researchers the opportunity to interactively explore antibodies clonal evolution. AncesTree displays interactive immunoglobulins (Ig) phylogenic tree, Ig related mutations and sequence alignments using additional information coming from specialized antibody tools (such as IMGT^®^). The GUI is a Java standalone application allowing interaction with Ig-tree that can run under Windows, Linux and Mac OS.

## Introduction

Development of Next Generation Sequencing (NGS) methodology and its use for high-throughput sequencing of the Adaptive Immune Receptor Repertoire (AIRR-seq) has provided unprecedented molecular insight into the complexity of the humoral adaptive immune response by generating Ig data sets of 100 million to billions of reads. Different computational methods have been developed to exploit and analyze these data (1). Retracing the antigen-driven evolution of Ig repertoires by inferring antibody evolution lineages is a powerful method to understand how vaccines or pathogens shape the humoral immune response (2–5). Indeed, Abs maturation is the result of clonal selection during B cell expansion. A clonal lineage is defined as immunoglobulin sequences originating from the same recombination event occurring between the V, D and J segments (6). Upon B cell receptor (BCR) engagement by a given antigen, somatic hypermutations (SHMs) events will generate a large BCR diversity, leading to antibodies with mutated Ig variable regions, thus forming a specific B-cell lineage that extends from the naive unmutated B-cells, to somatically hypermutated and class switched memory B or plasma-cells (7). Lineage tree building requires a common preprocessing step, where all sequences with identical V, J genes and CDR3 length (with a high CDR3 similarity) are grouped together (8–12). However, there is no consensus as to which phylogenetic method is optimal to infer the ancestral evolutionary relationships among Ig sequences (13, 14). Actually, several methods have been used, such as Levenshtein distance (LD), neighbor joining (NJ), maximum parsimony (MP), maximum likelihood (ML), and Bayesian inference (BEAST) (9, 15–17). DNA Maximum Likelihood program (Dnaml) of the PHYLIP package (18), is a ML method that has been originally developed for applications in ecology. It is also commonly used to infer B cell clonal lineages (19–24). Visualization of the phylogeny is performed using Dendroscope (25, 26). Currently there is no efficient bioinformatics tool allowing an interactive display of phylogenic tree inferred from Ig sequences. Here we developed AncesTree, a Dnaml Ig lineage tree visualizer that also integrates information coming from most used antibody bioinformatics tools: IMGT^®^ (27), Kabat numbering (28) and BASELINe (29). AncesTree enables users to interact with the tree generated by Dnaml via the GUI, which is a standalone application that is platform independent and only need JAVA JRE 12 or higher as prerequisite software installed.

### Design and implementation

The AncesTree workflow is presented in **Fig 1**, it consists of three different main steps: Input, Processing and Outputs.

**Figure 1.**
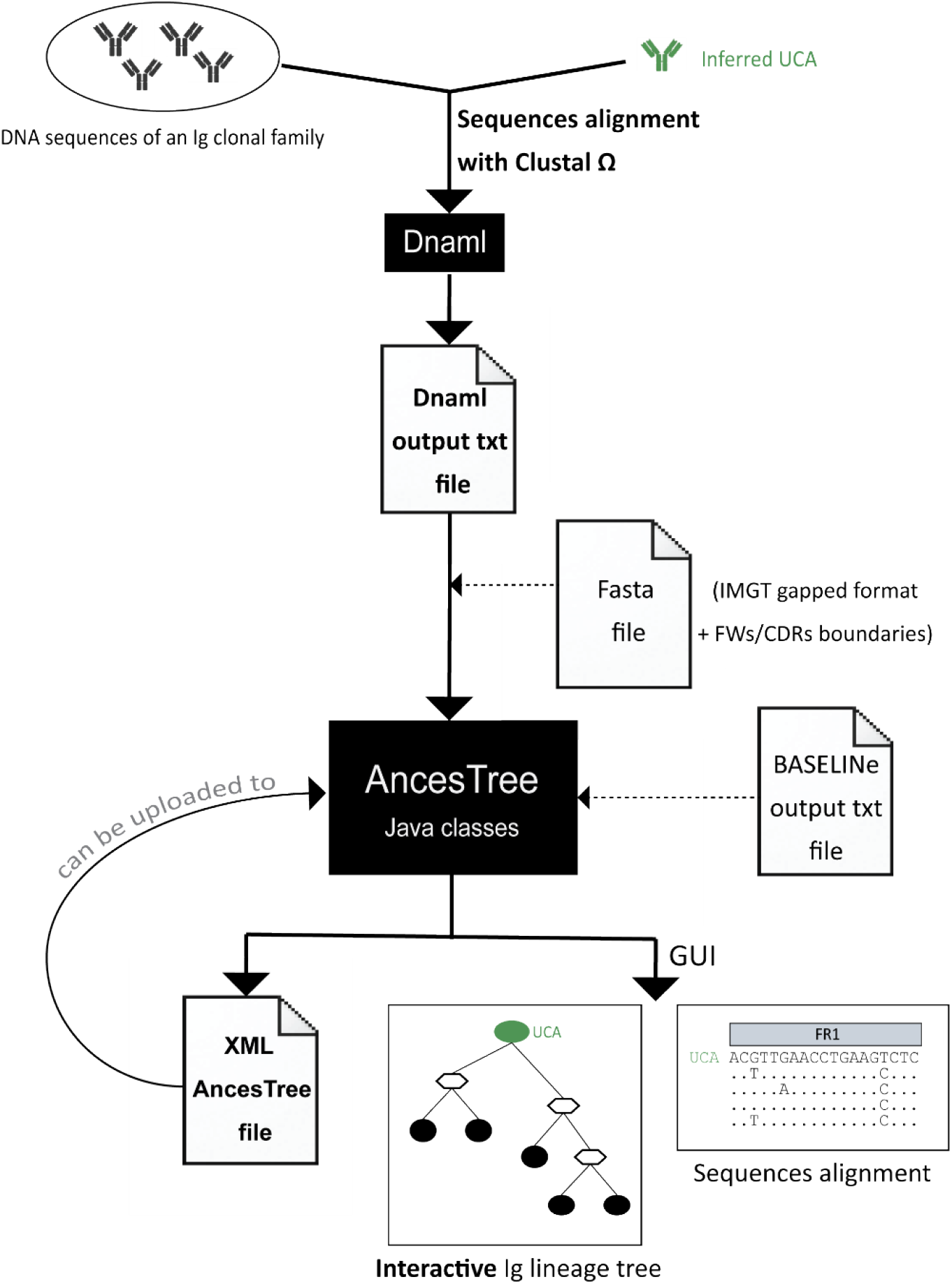
AncesTree workflow. DNA sequences of the variable region of an Ig clonal family of interest are aligned with Clustal Ω. Dnaml processes the sequences and generates a phylogenetic tree. The Dnaml output text file is then used as input for AncesTree. A fasta file with the UCA in gapped IMGT format can be provided (with the FWs and CDRs nucleotide positions in the fasta identifier). AncesTree processes the different inputs and reconstructs the phylogenic tree with all information related to Ig. BASELINe can be processed separately and its output saved in a text file and then uploaded into AncesTree. The tree is displayed in a GUI and an Extensible Markup Language (XML) file is produced (that could be used as direct input into AncesTree). Dashed arrows indicate optional features.

#### Input

The required input for AncesTree usage is the output text file generated by Dnaml. Optionally, a fasta file with data obtained from IMGT^®^ can also be used to have full AncesTree features.

A clonal family is composed of heavy (or light) V(D)J sequences and their related unmutated common ancestor (UCA). The UCA can be inferred with Antigen Receptor Probabilistic Parser (ARPP) UA Inference software (30) or Cloanalyst (31). Then, sequences are aligned with Clustal Ω (32) and the generated file in PHYLIP format can be provided for Dnaml. Dnaml is launched with the following settings: ‘O’ for the outgroup root with the number corresponding to the UCA position provided in the PHYLIP input text file and ‘5’ to reconstruct hypothetical sequences. The generated ‘outfile’ text file can be used as input for AncesTree.

To visualize the different frameworks (FW) and complementary-determining (CDR) regions that composed the Ig variable region, a fasta file can be uploaded. The user provides a fasta file containing the following information: the UCA V(D)J sequence in IMGT format including gaps, and the end positions of each region included in the fasta identifier (separated by a space). This information is easily retrieved using IMGT/V-QUEST (33) with the UCA nucleotide sequence as input.

#### Processing

AncesTree parses the Dnaml output file, and does not required a tree in Newick format. Indeed, the relationship between the different nodes of the tree is already stored, in addition to the sequence of each node, in the Dnaml output text file. The theoretical intermediate reconstructed sequences are renamed branch points (BPs) and in the case of ambiguous nucleotide notation (IUPAC nomenclature), AncesTree selects the nucleotide with the highest probability based on the Ig sequences retrieved after this BP. AncesTree has the ability to collapse a node if the sequences are identical, for example in the case of a theoretical BP correspond to an existing Ig. Moreover, AncesTree will also draw different nodes clustered together in the case of identical Ig sequences, thus providing a clear topology view of the tree.

#### Outputs

After running AncesTree, a sub-folder is automatically created in the ‘output’ folder, it uses the name of the Dnaml output file. The folder will contain all produced files such as a XML file that can be used for direct loading into the GUI.

AncesTree displays the processed tree in the main panel of the GUI **(Fig 2A)**. The number of nucleotide and amino acid mutations written on the edge between each node/sequence (with amino acid mutations shown in parenthesis) is clickable and enables the opening of a new window frame that displays the detailed location of each mutation **(Fig 2B)**. Of note, the color of the box around each mutated codon indicates whether the mutation is replacement (R) in red or silent (S) in green. This information is also available as R/S numbers under each region. The user can view the amino acid mutations, and have access by default to the Kabat numbering of the related amino acid position (without internet access, AncesTree will use the absolute position). To obtain the nucleotide or protein sequence of a node, the user can click on it **(Fig 2C)**. The user has also the possibility to enter the EC50 for the specified Ig. The sequence alignments (DNA or protein) are also accessible in a new frame via the ‘Menu’ button on the top **(Fig 2D)**. The alignment view is customizable: the sequences can be selected or deselected, as well as the different positions or regions. Different color modes can be chosen.

If the user is interested in a BASELINe analysis of its clonal family of interest, and if the optional input fasta file (with the UCA VDJ sequence including gaps) was provided, AncesTree generates automatically the fasta input file needed for this software (http://selection.med.yale.edu/baseline/). Once BASELINe is processed, its output can be loaded into AncesTree to have a nice graphic view of antigen-driven selection occurring for this particular clonal family. All generated graph can be exported in PNG or EPS format, the alignment can also be exported in a Tab-separated Values (TSV) file.

**Figure 2.**
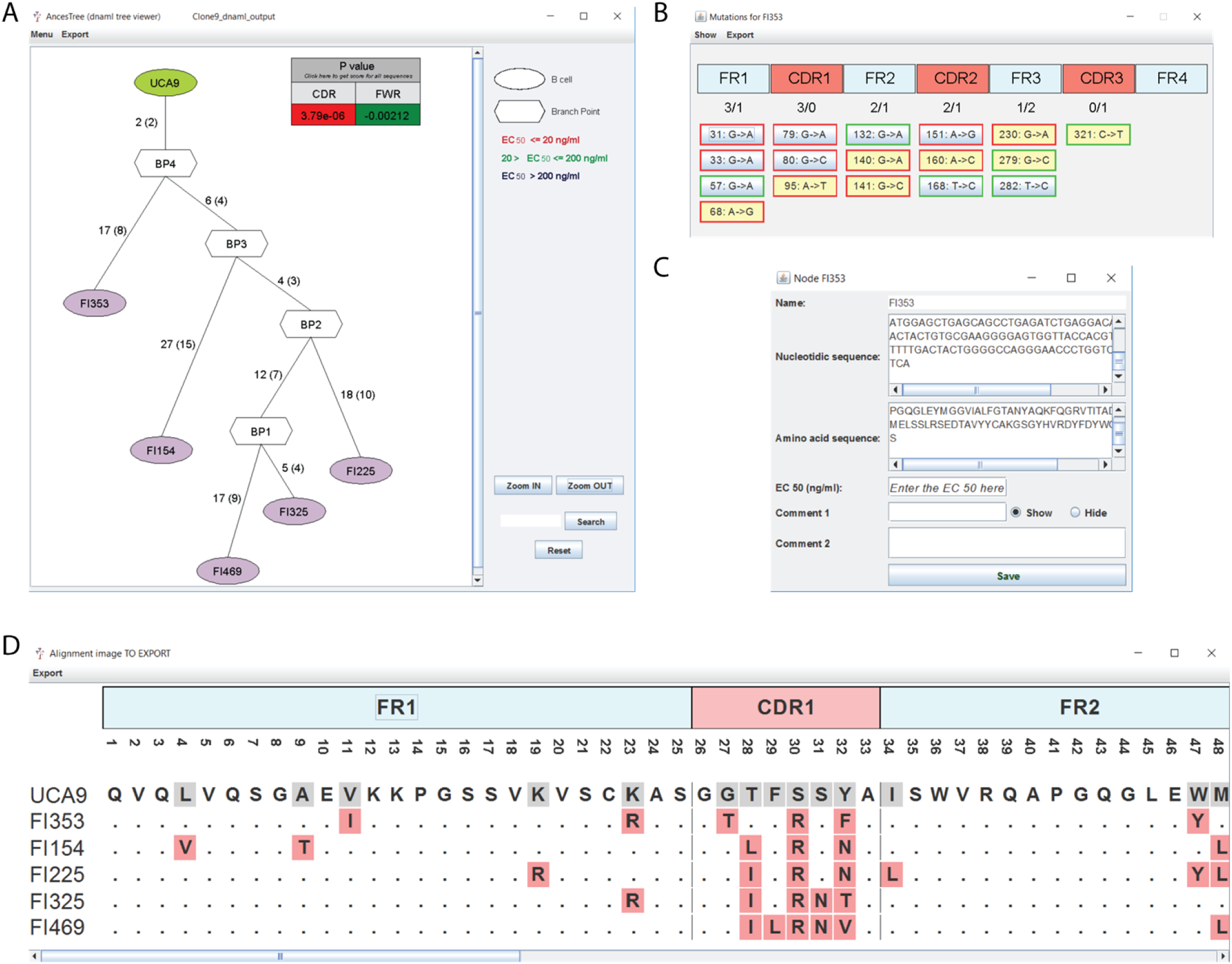
Snapshot of AncesTree GUI. (**A**) The tree generated by Dnaml is displayed in the main panel. The BASELINe analysis for the clonal family is displayed in the right upper corner. (**B**) The mutations between two nodes can be displayed in a separate window and they are positioned using IMGT^®^ sequence annotation. (**C**) The user can have access to each specific node to obtain the related sequences (DNA or protein) and add comments. (**D**) An alignment is generated with the UCA appearing in the first lane, and a ruler indicates the different regions that compose an Ig sequence.

## Results

To demonstrate the utility of AncesTree we analyzed a case study by performing the analysis of an Ig lineage tree targeting the fusion protein (F) of the Respiratory Syncytial Virus (RSV). RSV is an enveloped RNA virus belonging to the recently defined *Pneumoviridae* family (34). Infection of healthy adults by RSV typically results in mild respiratory symptoms. However, viral infection of infants and older adults, accounts for a substantial hospitalization burden in both age groups (35). Indeed, RSV infection is the second cause of infant mortality worldwide after malaria (36). Understanding the immunological basis for the development of potent neutralizing antibodies is a key step for the development of an effective vaccine for RSV.

### Case study: Exploration of Ig lineage targeting the Fusion protein of the Respiratory Syncytial Virus (F-RSV)

To demonstrate the practical use of AncesTree, we re-analyzed an Ig dataset generated post infection by Respiratory Syncytial Viral infection (HRSV). The dataset was collected by isolating antibodies direct against the RSV F protein, a class I fusion protein mediating viral entry into host cells (37). The Ig sequences were clustered by grouping antibodies sharing the same VH and VL gene usage, HCDR3 length and identity (at least 85% for HCDR3). Among the clusters generated, we chose Igs targeting the antigenic site V of RSV F located near amino acid 447 between the α3 helix and β3/β4 hairpin of F-RSV in prefusion **(Fig 3A)**. About 70% of the mAbs targeting this site use the same VH and VL germline pair (VH1–18 and VK2–30) (37–39). We identified an Ig family of interest containing potent neutralizers targeting site V with one outlier, the mAb ADI-14576, being less potent and with a 10-fold decrease in binding affinity (**Fig 3B**). We used Dnaml to generate VH sequences phylogenic tree and launched AncesTree to analyze and interact with the produced phylogenic tree (**Fig 4A**). The EC50 (ng/ml) related to the neutralization assay against RSV subtype A are reported in each node (of note, EC50 against subtype B are in the same range for each Ig). Surprisingly, a common mutation 92:G->A (kabat position 31: S ->N) is shared between all the Igs, except for ADI-14576 that does not share this mutation. The alignment of the Ig protein sequences highlights clearly this shared mutation (**Fig 4B**). A result suggesting that ADI-14576 underwent less affinity maturation and therefore diverges from all the other family members. Interestingly, the 31:S->N mutation is located in the HCDR1 and asparagine residues are often involved in protein binding sites. It is tempting to speculate that the Serine to Asparagine substitution is in part responsible for the higher potency and binding titer of the antibodies.

**Figure 3.**
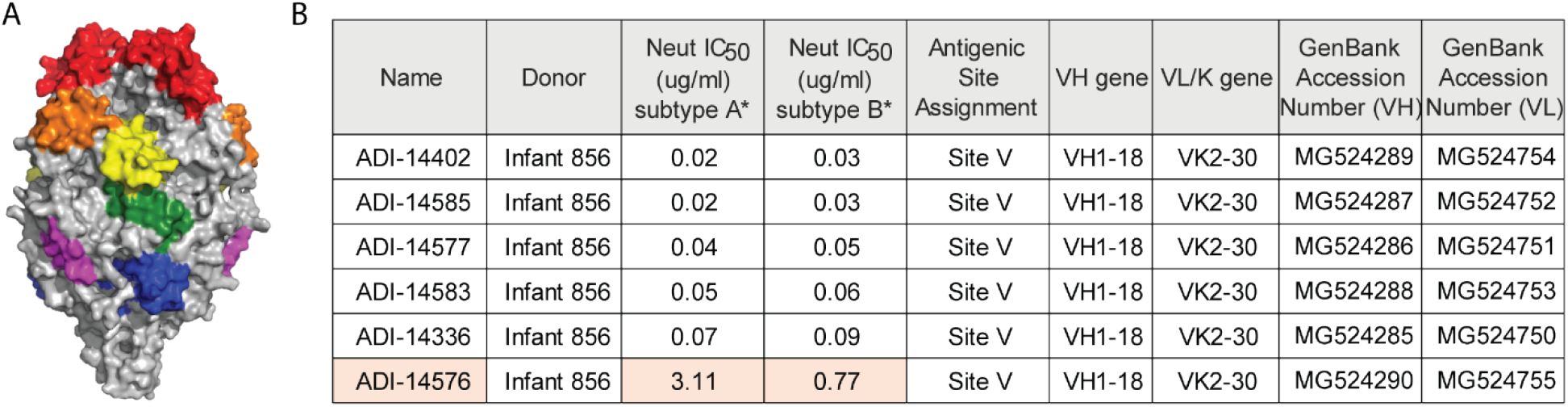
Clonal family against RSV-F protein antigenic site V. (**A**) Shown is the prefusion conformation of RSV F trimer. The antigenic sites are colored, site Ø (red), I (blue), II (yellow), III (green), IV (purple) and V (orange). Representation was done using PDB ID 4mmu (40) and prepared using PyMOL software (The PyMOL Molecular Graphics System, Version 4.5 Schrödinger, LLC). (**B**) Table showing the different characteristic of a mAbs clonal family isolated from an infant (≥ 6 months) after RSV infection. The Igs neutralization titers are shown as well as their related Germline annotations. ADI-14576 is highlighted because of is lower neutralization value in comparison to the other mAbs of the same clonal family.

**Figure 4.**
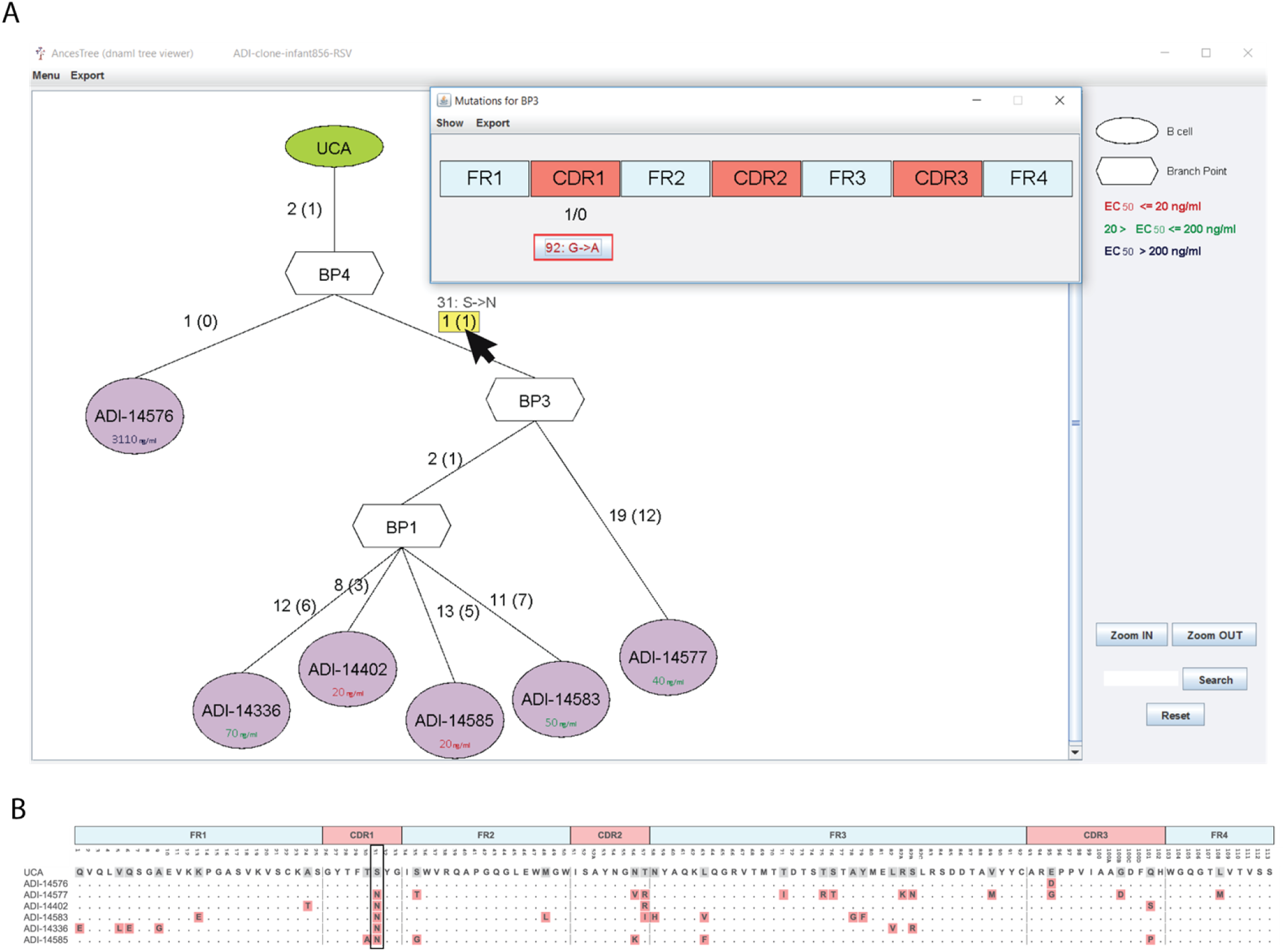
Phylogenetic analysis of the VH chain of a clonal family RSV-F specific. (**A**) Phylogenic tree displayed in AncesTree where the user clicked on the shared mutation for all Igs below BP3 node (31: S->N). (**B**) Protein alignment of the different Ig sequences, the mutation 31: S->N is boxed.

### Concluding remarks

To summarize, we developed an intuitive, easy and interactive GUI allowing the visualization and exploration of antibody clonal evolution. Our application is open access and only needs the file produced by Dnaml and restricted information specific to antibody sequence analysis.

## Availability

AncesTree is open-source software implemented in Java and freely available from https://bitbucket.org/mathildefog/ancestree. Documentation for installation and user tutorial are provided.

## Authors’ contributions

MF developed the application and performed the analyses. LPa, AL, DC and LPe participated in the design of the application. MF and LPe wrote the paper. All authors read and approved the final manuscript.

## Competing interests

The authors declare that they have no competing interests.

## Funding

This research did not receive any specific grant from funding agencies in the public, commercial, or not-for-profit sectors.

## Acknowledgements

The authors acknowledge present and past members of the Lanzavecchia’s group for comments and feedback on the software.

